# Unravelling the Interplay of Nitrogen Nutrition and the *Botrytis cinerea* pectin lyase BcPNL1 in Modulating *Arabidopsis thaliana* Susceptibility

**DOI:** 10.1101/2024.03.06.583732

**Authors:** Antoine Davière, Aline Voxeur, Sylvie Jolivet, Samantha Vernhettes, Marie-Christine Soulié, Mathilde Fagard

## Abstract

In this study, we investigated the intricate interplay between nitrogen nutrition, and the dynamics of pectin degradation during plant-pathogen interactions, using *Arabidopsis thaliana* and *Botrytis cinerea* as a model pathosystem. Our findings revealed a noteworthy impact of nitrogen availability on the pectin degrading activity of the *B. cinerea* pectin lyase 1 (PNL) for which the mutant strains presented a reduced pathogenicity restored by complementation. More precisely, infected nitrogen-sufficient (high N) plants exhibited an increased release of PNL-derived oligogalacturonides compared to infected nitrogen-deficient (low N) plants. This correlated with an elevated expression of jasmonic acid repressor genes in high N plants, rendering them more susceptible to *B. cinerea*. Using *ΔBcpnl1* deletion mutants, we demonstrated that the increased production of BcPNL1 -derived oligogalacturonides under high N conditions was responsible for the increased expression of jasmonic acid repressor genes, significantly contributing to the higher susceptibility of high N plants to *B. cinerea.* In conclusion, we demonstrated that BcPNL1 is a major pathogenicity factor during *B. cinerea* interaction that is affected by the plant’s N nutrition conditions.

## Introduction

The ascomycete *Botrytis cinerea* is a widely spread necrotrophic fungus responsible for the grey mold disease in over 1000 plant species (Fillinger & Elad, 2015). As a necrotrophic pathogen, its infectious process relies on several synergistic molecular mechanisms, involving the secretion of a vast array of molecules that include cell wall-degrading enzymes, toxins, reactive oxygen species, peptidases, and acidifying compounds, which have been reported to contribute to its pathogenicity (Cheung et al., 2020).

The availability of the sequenced genome of this fungus (Amselem et al., 2011), has greatly facilitated genetic studies and the identification of putative pathogenicity factors. However, the high degree of functional redundancy and compensation phenomena in this fungus has made it challenging to pinpoint the roles of individual *B. cinerea* genes in pathogenicity. Consequently, only a few genes have been described as having a major role in pathogenicity. Among them, some transcription factors and kinases have been identified, such as sucrose non-fermenting protein kinase 1 (BcSNF1) and the pH regulator BcPACC (Lengyel et al., 2022; Rascle et al., 2018). Furthermore, chitin synthases like Bcchs3a are also important pathogenicity factors, influencing fungal adhesion to the host and activating plant defense mechanisms (Arbelet et al., 2010). In addition to these, several secreted factors, such as the botrydial and botcinic acid toxins whose production involve the *BcBOT* and *BcBOA* gene clusters, as well as cell death-inducing proteins like BcXIG1 and BcCRH1, play significant roles in pathogenicity (Bi et al., 2021; Dalmais et al., 2011; Zhu et al., 2017). However, recent papers have raised concerns about studying pathogenicity factors exclusively in optimum conditions since the plant -pathogen interaction is strongly impacted by abiotic factors (Farjad et al., 2021; Zarattini et al., 2021). Indeed for *B. cinerea*, we were able in a previous work to identify undescribed pathogenicity factors by varying plant nitrogen supply which significantly affect the interaction (Soulie et al., 2020).

During infection, *B. cinerea* degrades the cell wall of the plants it infects. To achieve this, the pathogen releases numerous enzymes, with the most abundant belonging to the pectinase family (Espino et al., 2010). Pectinases enable the pathogen to penetrate and spread within host tissues by degrading pectins, one of the major components of the plant cell wall barrier. The primary content of pectin, homogalacturonan (HGA), consists of α-1,4-linked residues of D-galacturonic acid (GalA), which can be methyl (Me)- and/or acetyl-esterified (Ac). The degradation of HGA by pectinases leads to the production of oligogalacturonides (OG) with varying degrees of polymerization (DP) and methyl-acetylesterification. Among pectinases, pectin lyases (PNLs) are likely the primary OG-producing pectinases of *B. cinerea* during early infection events in tomato fruit (An et al., 2005) and *Arabidopsis thaliana* leaves (Voxeur et al., 2019). PNLs employ β-elimination to degrade highly methylesterified pectin, resulting in highly methyl- and acetylesterified OGs harboring an unsaturated bond at the non-reducing end (Yadav et al., 2009). Subsequently, esterases, including pectin methylesterases (PME) and pectin acetylesterases (PAE), deesterify the large methyl- and acetylesterified oligosaccharides, allowing depolymerases — which degrade low and non-methylesterified pectin — to further break down these large oligosaccharides into smaller ones. These depolymerases are categorized into two groups: hydrolases with polygalacturonases (Endo-PGs and Exo-PGs) such as BcPG1 and BcPG2 which are crucial pathogenicity factors (Ten have et al., 1998; Kars et al., 2005) and collectively represent 20% of the total protein content in the early secretome of *B. cinerea* (Espino et al., 2010) and lyases with pectate lyases (Endo-PLs and Exo-PLs).

Plants perceive the oligogalacturonides (OGs) produced by these pectinases as signals of damaged or modified self, enabling them to act as Damage-Associated Molecular Patterns (DAMPs) in Pattern-Triggered Immunity (PTI) (Pontiggia et al., 2020). This signaling pathway triggers several modifications in plant cells, leading to the accumulation of defense proteins and the production of hormones that enhance the defense reaction. Recently, Davidsson et al. in 2017 showed that short OGs (DP3) can efficiently trigger hormonal immunity mediated by jasmonic acid (JA) and salicylic acid (SA). These short OGs resulting from PG activity effectively protected *A. thaliana* from *Pectobacterium carotovorum* infection and are detected in *A. thaliana* leavesinfected by *B. cinerea* (Voxeur et al., 2019). Interestingly, research has also shown that OG oxidases (OGOX) secreted by the plant itself can inactivate OG-induced defense, thus preventing excessive plant cell death and infection during *B. cinerea* infection (Benedetti et al., 2018). The hydrogen peroxide (H_2_O_2_) produced during OG oxidation mediates the activation of other defense processes, such as lignification or the inactivation of auxins, through the action of class III plant peroxidases (Scortica et al., 2022). Finally, early reports on OG activity demonstrated that long-chain OGs as a result of PG activity on fully demethylesterified HGA (DP10-15) serve as defense elicitors and can induce phytoalexin production (Ferrari et al., 2007; Hahn et al., 1981). However, we failed to detect these long-chain OGs in *A. thaliana* leaves infected by *B. cinerea* (Voxeur et al., 2019) as well as in infected tomato fruit (An et al., 2005). Instead, we detected unsaturated methylesterified OGs with long DP capable of suppressing JA-mediated defense activation by up-regulating the pool of transcripts encoding negative regulators of JA signaling (Voxeur et al., 2019). However, the subsequent degradation of these OGs by PMEs and PGs leads notably to the production of GalA_4_MeAc-H_2_O, which, in the end, strongly elicits JA-related plant defense.

To havea better insight on *B. cinerea* strategiesto escape plant defense, we investigated the role of BcPNLs in pathogenicity in a context of environmental modulation of the plant-pathogen interaction using nitrogen fertilization. Our research provides evidence that nitrogen fertilization impacts *A. thaliana* pectin sensitivity to PNLs and that BcPNL1 contributes to the factors linking disease susceptibility and nitrogen fertilization. More precisely, by conducting mutant and complementation analyses, our findings demonstrate that BcPNL1 plays a paramount role in *B. cinerea* pathogenicity and confirm the involvement of BcPNL1 in producing unsaturated methylesterified OGs with long DP capable of suppressing plant JA-mediated defense activation.

## Results

### Host nitrogen nutrition impacts pectin degradability during A. thaliana/B. cinerea interaction

First, we assessed if the host nitrogen nutrition impacts the OG turnover during infection by analysing the OG produced in presence of the WT strain B05.10 on plants grown either with 0.5 mM (low N) or 10 mM NO_3_^-^ (high N) which correspond physiologically to nitrogen-deficient or -sufficient regimes respectively (Soulie et al., 2020). To do so, we incubated plant leaves in a liquid medium containing fungal spores of the WT strain B05.10 and subsequently analyzed the cell wall breakdown products released into the medium using high-performance size-exclusion chromatography (HP-SEC) in conjunction with a high-resolution mass spectrometry (HRMS) (Voxeur et al., 2019). Interestingly, we found a higher quantity of PNL products of DP6, 7 and 9 at high N compared to low N leading to an increased PNL products proportion and a decreased PG products proportion (Figure 1A).

We next assessed if host nitrogen nutrition could impact fungal pectinase gene expression. As depicted in Figure 1B, at 1 day post-infection (dpi), *BcPG1,2* and *BcPNL1* stand out as the most highly expressed genes within the *BcPG* and *BcPNL* gene families (Figure S1), respectively, at both high and low N. Moreover, we observed that at 1 dpi, the expressions of *BcPNL1, 2* were decreased at high N (Figure 1B) whereas we detected more PNL products. It suggested that nitrogen nutrition impacts host pectin structure that in turn impacts OG production rather than the fungal pectin degradation machinery.

Since PNL products of DP>4 have been shown to up-regulate *JOX* and *JAZ* genes (Voxeur et al., 2019) respectively involved in repression of the JA pathway by inactivation of JA through hydroxylation (Caarls et al., 2017; Smirnova et al., 2017) and repression of JA-induced genes (Zhang et al., 2015), we searched for differences in nitrogen-dependent *JAZ* and *JOX* regulation during infection. We found a significant higher expression of *AtJAZ*3,4 and *AtJOX3* in high N compared to low N at 1 dpi which was consistent with the higher production of PNL products upon high N plant infection (Figure 1C). Accordingly, the JA marker gene *AtPDF1.2* was less expressed in high N conditions which was also previously observed in Soulie et al., 2020. Since SA and JA defence signalling pathways are mutually antagonistic, we also searched for SA marker genes. Upon infection, we observed at high N a higher expression level for *AtPR1* and the transcription factor *AtWRKY70* which is activated by SA and repressed by JA (Li et al., 2004).

**Figure 1:**
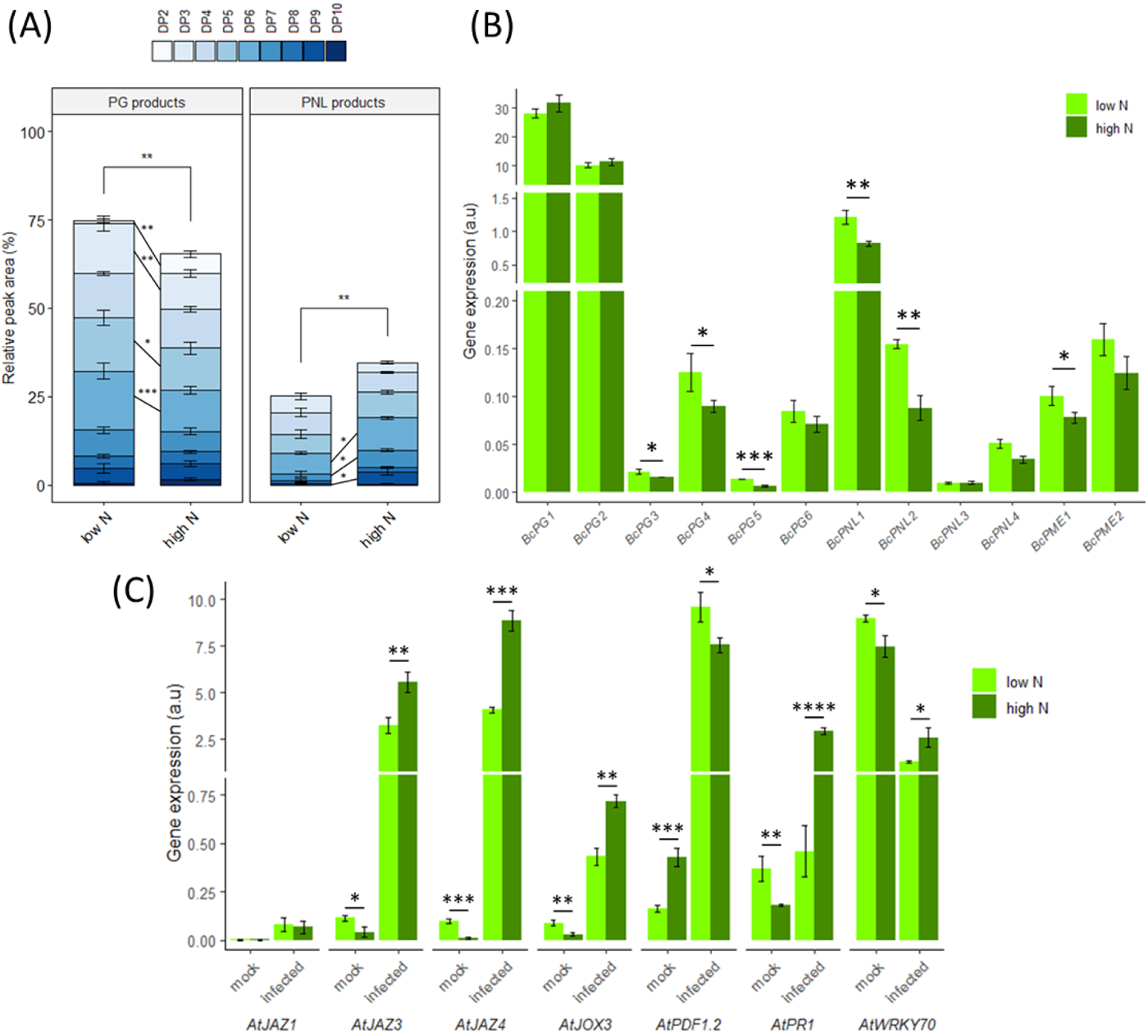
Impact of host nitrate nutrition on Botrytis cinerea pectinolytic capacities and plant defense regulation. (A) LC-MS quantitative analysis of OG production after incubation of spores from the WT strain of the fungus for 15 h with A. thaliana Col-0 leaves from plants cultivated at low N or high N. (B,C) Transcript accumulation during infection in low N and high N conditions at 1 day post infection are represented for (B) fungal pectinase genes and (C) plant defense genes. Fungal gene targets were normalized with BcActA whereas plant gene targets were normalized with AtUBI4. Similar results were obtained using BcUBI on the fungus side or AtAPT1 on the plant side as reference genes for normalization. Three independent experiments were conducted with similar results. Data are expressed as mean normalized expression in arbitrary units (a.u.) and are the means of triplicates (±SE). Statistical differences represented between low N and high N are the results of a two -way ANOVA followed by a Fisher’s LSD test in (A) and a two sample t -test in (B) and (C): *p<0.05, **p<0.01, ***p<0.001, ****p<0.0001.

### Targeted gene deletion of the BcPNL1 gene in B. cinerea leads to reduced pathogenicity at high N and low N

Next, to draw a clear connection between the higher accumulation of PNL products and the downregulation of the JA defense signalling pathway in high N compared to low N plants, we investigated the function of *BcPNL1* which stands out as the most highly expressed gene within the *BcPNL* gene family as depicted in Figure 1B. To investigate its function, we replaced *BcPNL1* coding region with a deletion cassette containing the fenhexamid resistance gene, which was amplified from pTEL-fenh (Leisen et al., 2020). We employed PEG/CaCl2-mediated transformation of *B. cinerea* protoplasts, in conjunction with the recently described CRISPR-Cas9 system, utilizing the ribonucleoprotein complex direct delivery strategy (Leisen et al., 2020, as shown in Figure 2A). *In vitro*, we selected fenhexamid-resistant transformants.

The absence of the wild-type *BcPNL1* gene and the correct integration of the cassette in each isolate were determined through PCR and confirmed via Southern blot using a DIG-labeled probe. This approach also allowed us to verify the absence of ectopic integration of the cassette in the genome (see Figure 2B). Figure 2B demonstrates that out of 20 resistant transformants, we successfully identified three homokaryotic mutant strains, designated *ΔBcpnl1.1*, *ΔBcpnl1.2*, and *ΔBcpnl1.3*. Others, such as the *ΔBcpnl1.4* transformant, were found to be heterokaryotic. Nevertheless, all tested strains were devoid of ectopic integration, confirming the precision of the CRISPR-Cas9 system (as shown in Figure 2C). Subsequently, we selected the mutant strains *ΔBcpnl1.1, ΔBcpnl1.2,* and *ΔBcpnl1.3* for further phenotypic characterization.

**Figure 2:**
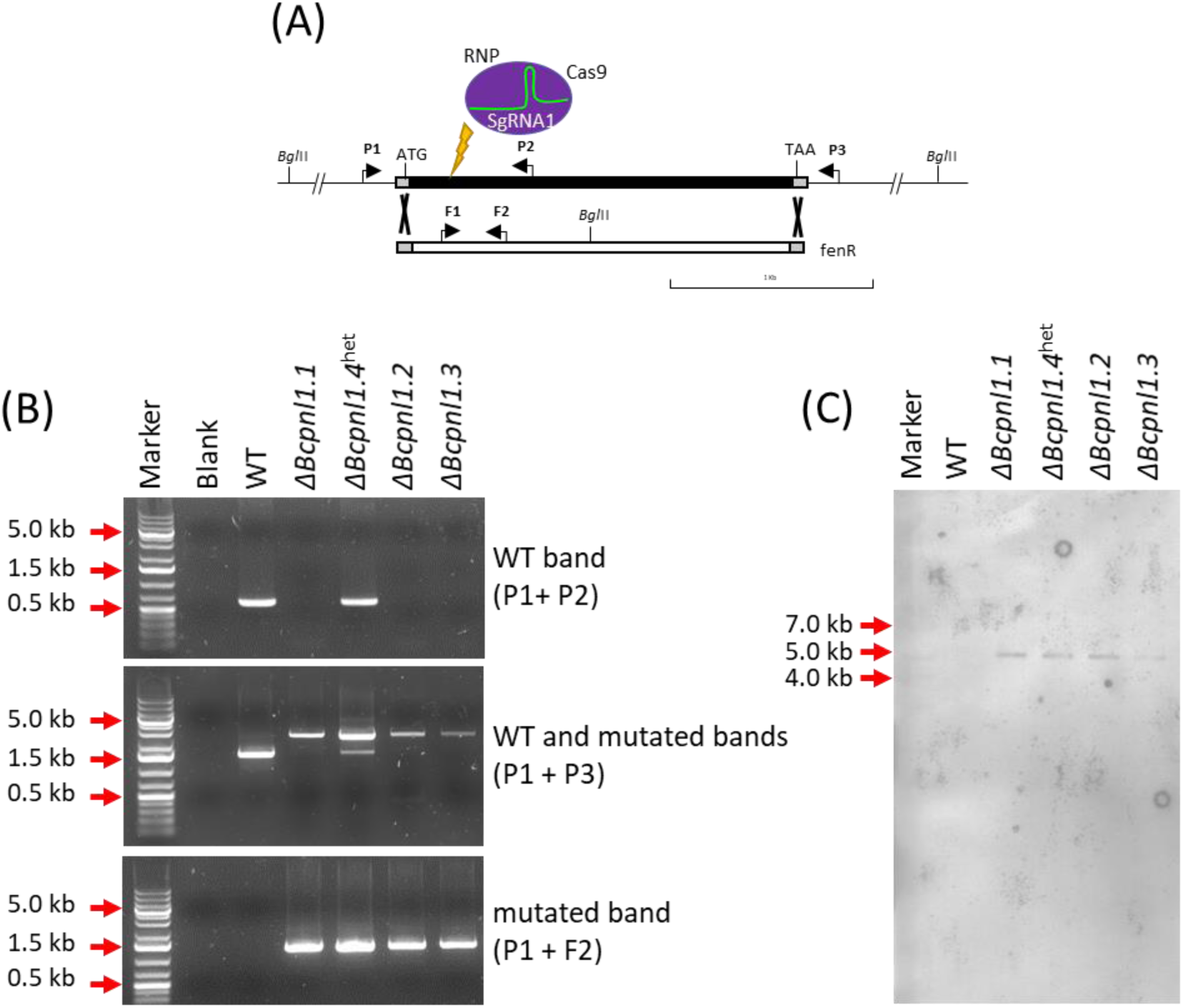
Targeted gene replacement of BcPNL1 in Botrytis cinerea. (A) Representation of the deletion strategy used to obtain Bcpnl1 mutants (arrows indicate primers P1, P2 and P3 used for diagnostic PCR shown in panel (B) and Southern in panel (C)). (B) Diagnostic PCR showing successfull integration of the fenR cassette and nuclear purity of the ΔBcpnl1.1, ΔBcpnl1.2 and ΔBcpnl1.3 mutants whereas ΔBcpnl1.4 is heterokaryotic. (C) Southern hybridation of genomic DNA. In order to detect multiple integration events, genomic DNA was digested by BglII and hybridized with a fenR cassette probe obtained by PCR with F1 and F2 primers. Only one band of the expected size was found in all transformants.

First, we investigated whether the deletion of *BcPNL1* would affect the growth of the fungus *in vitro* when cultivated in solid medium in the presence of pectin. Although we observed slight differences between the mutant strains, we did not observe any significant impact of *BcPNL1* deletion on *in vitro* spore production on PDA or growth with Czapeck minimal medium supplemented with glucose or highly methylesterified pectin as the sole carbon sources (Figure S2A). However, the *ΔBcpnl1* mutants did exhibit a delay in spore germination on onion epidermis compared to the wild-type B05.10 strain, although penetration was still observed at the 24-hour time point as for the wild-type (Figure 3B).

Next, we assessed whether the deletion of *BcPNL1* could affect pathogenicity on low and high N plants. The *ΔBcpnl1* mutants exhibited a substantial reduction in pathogenicity on both Columbia (Col-0) and Wassilewskija (Ws) *A. thaliana* leaves (Figure 4A-B) under both nitrogen nutrition conditions at 2 dpi. qPCR quantification of fungal DNA during the infection of *A. thaliana* Col-0 leaves confirmed the reduced pathogenicity of these mutants (Figure 4C). Furthermore, we observed an increased susceptibility of high N plants infected either by B05.10 or *ΔBcpnl1* mutants suggesting that BcPNL1 pectinolytic activity does not fully contribute to the impact of host nitrogen nutrition on disease severity. We then confirmed that the role of BcPNL1 in pathogenicity is not restricted to *A. thaliana* infection since tomato leaves infected with the *ΔBcpnl1* mutants also presented smaller lesions compared to B05.10 (Figure 4D-E).

Lastly, we complemented the *ΔBcpnl1.1* mutant with the *BcPNL1* gene either under the control of the native promoter (1.5kb upstream of the ORF) or the strong *oliC* promoter from *Aspergillus nidulans*, resulting in the *ΔBcpnl1.1c1* and the *ΔBcpnl1.1c2* strains respectively (Figure 5A-B). The complementation cassettes were integrated at the same intergenic region of chromosome 3 using CRISPR-Cas9 as described previously (Jeblick et al., 2023). With both complemented strains we observed a partial restoration of the pathogenicity on Col0 *A. thaliana* leaves (Figure 5C). RT -qPCR analysis of the complemented lines revealed that the level of *BcPNL1* expression was only partially restored compared to the B05.10 strain. This may explain why we did not achieve a complete restoration of the pathogenicity (Figure 5D).

**Figure 4:**
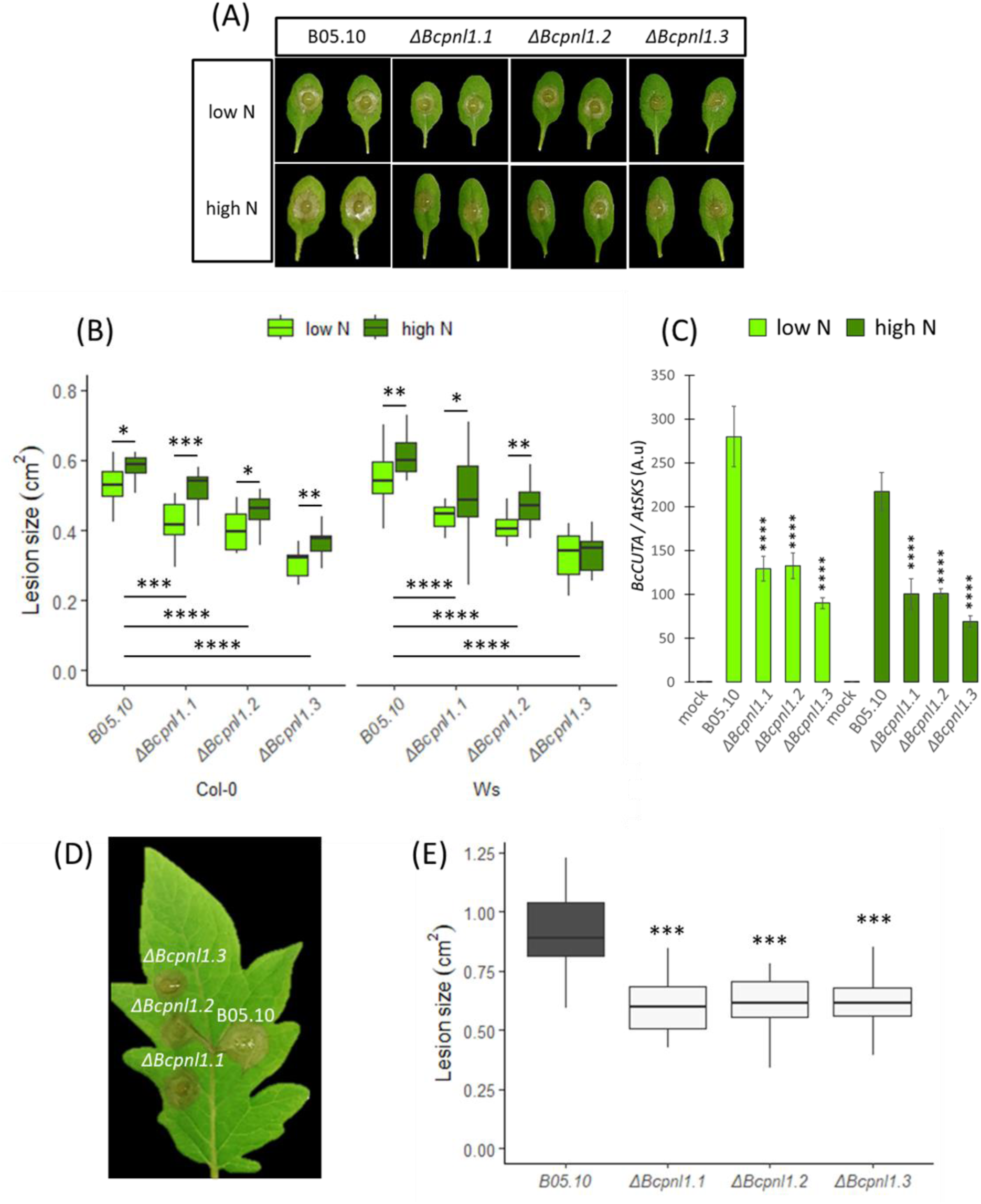
Pathogenicity test on detached leaves with the B05.10 reference strain of Botrytis cinerea and the ΔBcpnl1 mutant strains. (A) Columbia A. thaliana leaves at 2dpi. (B) Plot and statistical analysis of the lesions on Columbia and Wassilewskija A. thaliana leaves. Plants were either grown with low or high nitrate supply (low N, high N). (C) Fungal DNA quantification of B05.10 and the ΔBcpnl1 mutants strains of B. cinerea during infection of A. thaliana Col-0 plants. Data are expressed as mean quantification of the fungal BcCUTA gene over the plant AtSKS gene in arbitrary units (a.u.) and are the means of triplicates (±SE). (D) S. lycopersicum leaves, 2 dpi. (E) Plot and statistical analysis of the lesions on tomato leaves at 2dpi. All the tomato plants were grown with the same N supply (10 mM NO_3_^-^). In each condition, two sample t-test was conducted between the B05.10 strain and the mutant strains: *p<0.05,**p<0.01,***p<0.001. All infection experiments were done with at least 15 leaves and repeated three times with similar results.

**Figure 5:**
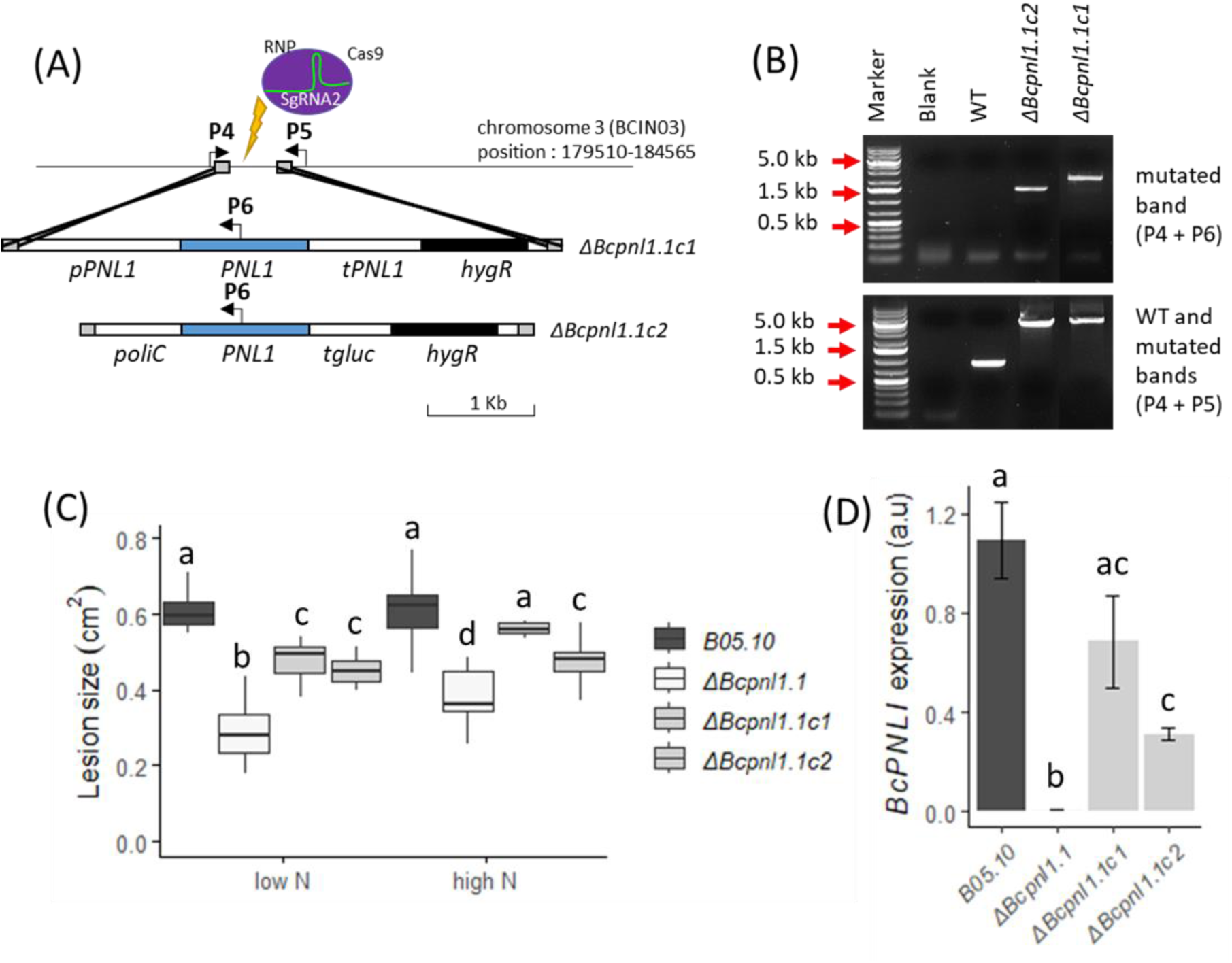
Complementation of the ΔBcpnl1.1 mutant partially restored the in planta phenotype. (A) Targeted insertion of the BcPNL1 complementation cassette in the ΔBcpnl1.1 mutant strain of B. cinerea. Arrows indicate primers used for diagnostic PCR shown in panel (B). (B) Diagnostic PCR showing successful integration of the complementation cassette and nuclear purity of the ΔBcpnl1.1c1 and ΔBcpnl1.1c2 strains. (C) The plot represents the lesions on A. thaliana detached leaves at 2dpi with B05.10, the ΔBcpnl1.1 mutant and the two complemented strains ΔBcpnl1.1c1 and ΔBcpnl1.1c2. (D) Transcript accumulation of the BcPNL1 gene at 1dpi on A. thaliana Col-0 leaves (high N) with the different strains of B. cinerea. BcActA was used for normalization of BcPNL1 expression. Letters represent the results of a two-sample t-test, p<0.05. All infection experiments were done with at least 15 leaves and repeated three times with similar results.

### ΔBcpnl1 mutants produced less PNL and PG products at high N and low N

Next, we investigated the impact of *BcPNL1* deletion on cell wall degradation. The analysis of pectin degradation products in the presence of the *ΔBcpnl1* mutants at 15 hours post-inoculation (hpi) revealed on one hand a drastically reduced production of PNL derived OGs compared to the WT strain, in both high N and low N plants (Figure 6A), confirming the importance of BcPNL1 on the total fungal PNL activity. On the second hand, PG products were also strongly reduced upon infection with the mutants, indicating that the PNL activity is essential upon infection to regulate PG and PME activity. In order to assess if BcPNL1 activity could regulate the other pectinases at the transcriptional level or if the decrease of PG products only results from the inability of PMEs or PGs to have access to their substrate without prior PNL activity, we assessed the level of expression of pectinase genes in *ΔBcpnl1* mutants at 1 dpi (Figure 6B). Interestingly, all the tested pectinase genes were downregulated in the *ΔBcpnl1* mutants compared to the WT strain in both N conditions.

Next, we performed an OG analysis at 18 hpi on plants grown either on high N or low N and infected by *ΔBcpnl1.1c1* and the *ΔBcpnl1.1c2* strains. At this later time point than previous analysis, high DP OGs couldn’t be detected but we could still observe that the production of OGs was partially restored with the complemented strains compare to the *ΔBcpnl1.1* mutant (Figure 6C). More precisely, at 18 hpi, we observed a restoration of DP4 and DP5 PNL products in low N plants but still less PNL products in high N plants infected by the complemented strains suggesting that the level of complementation was high enough to restore the full OG turnover speed (degradation of high DP OGs to produce shorter ones) during infection of low N but not of high N plants. Furthermore, PG products were still less abundant in high N plants infected by *ΔBcpnl1.1c2* strain confirming that the PNL is essential prior to PG activity (Figure 6C)

**Figure 6:**
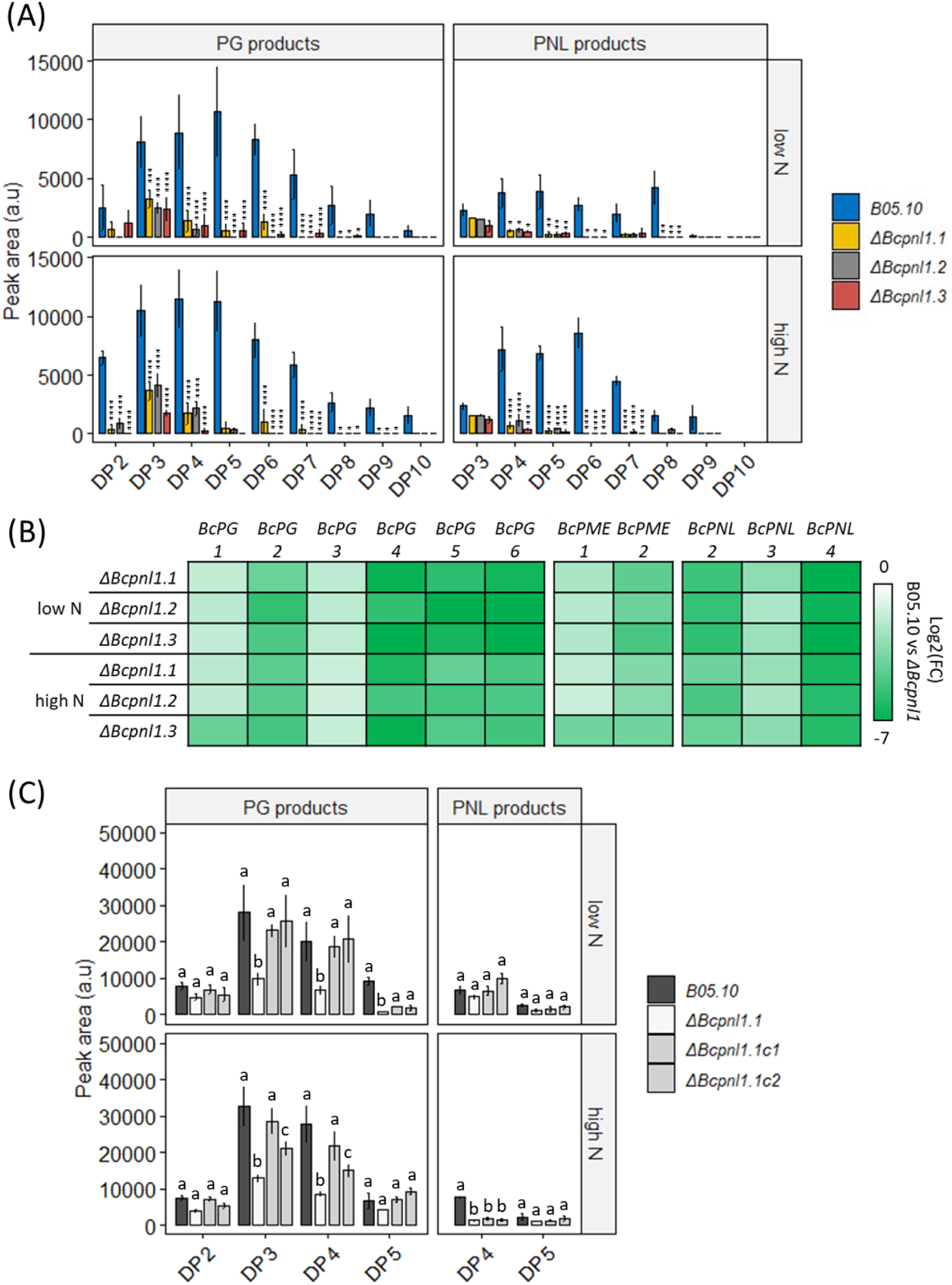
Impact of BcPNL1 mutation on Botrytis cinerea pectinolytic capacities. (A) LC-MS analysis of OG production from leaves of A. thaliana plants ( cultivated at low N or high N) incubated for 15 h in liquid medium supplemented with spores of the wild-type and mutant strains of B. cinerea . A two-way ANOVA test was performed followed by a Fisher’s LSD test between wild type and mutant strains: *p<0.05, **p<0.01, ***p<0.001. (B) The log2 fold change of transcript accumulation during infection in low N and high N conditions at 1 dpi are represented for fungal pectinase genes. Fungal gene targets were normalized with BcActA whereas plant gene targets were normalized with AtUBI4. Similar results were obtained using BcUBI for normalisation. Three independent experiments were conducted with similar results. Data are expressed as mean normalized expression in arbitrary units (a.u.) and are the means of triplicates (±SD). Statistical differences represented between low N and high N are the results of a two-way ANOVA followed by a Fisher’s LSD test in (A) and a two sample t-test in (B) and (C) : *p<0.05, **p<0.01, ***p<0.001 . (C) LC-MS analysis of OG production from leaves of A. thaliana plants (cultivated at low N or high N) incubated for 18 h in liquid medium supplemented with spores of the WT, mutant and complemented strains of B. cinerea. Letters represent the results of a two-way ANOVA test followed by a Fisher’s LSD test between strain for each DP group. DP: degree of polymerisation.

### Infection with ΔBcpnl1 mutants triggers reduced cell death and oxidative burst

To characterize the plant response to the hypovirulent *ΔBcpnl1* mutants, we analyzed the oxidative burst by measuring H_2_O_2_ production using a 3,3’-diaminobenzidine (DAB) staining at an early time point after inoculation. At 1 dpi, high and low N Col-0 leaves infected with B05.10 strain exhibited a strong DAB staining at the inoculation site and its vicinity, while leaves infected with the *ΔBcpnl1* mutants showed only slight staining at the inoculation site (Figure 7). We also assessed plant cell death upon infection using trypan blue (TB) staining. At the same time point (1 dpi), TB staining revealed fungal growth as well as small, darker spots indicating plant cell death, but this was observed only for the B05.10 strain at high and low N (Figure 7B, 7C). These results indicate that at this early time point, the plant had responded to the pathogenic B05.10 strain but not to the mutants. Taken together, these results suggest a lower aggressiveness of the *ΔBcpnl1* mutants compared to B05.10 at high and low N, consistent with the findings from the pathogenicity tests and OG analysis.

**Figure 7:**
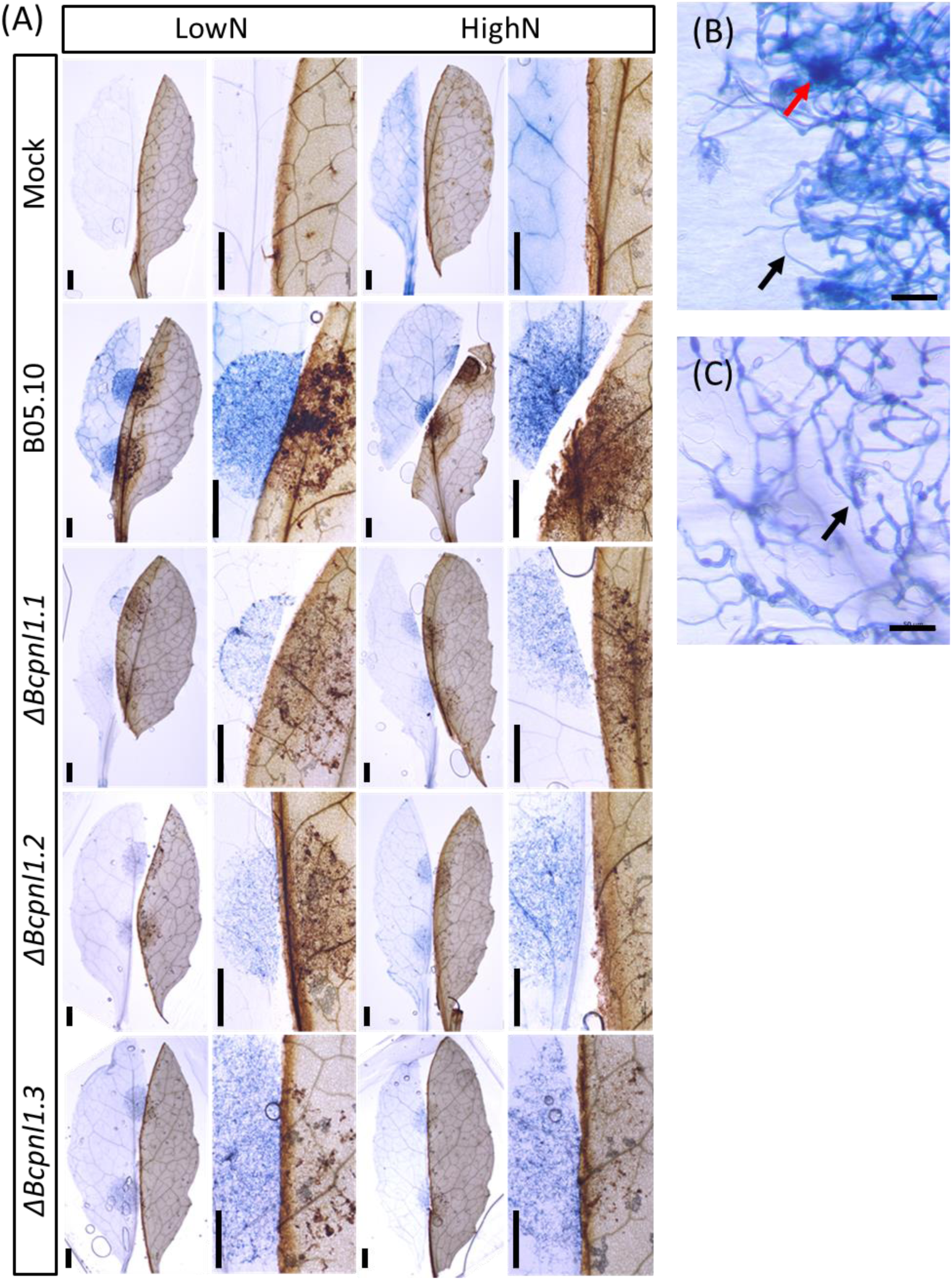
Cell death and oxidative burst reaction during infection with Botrytis cinerea. Detached leaves of A. thaliana were infected with drops either only with ½ PDB (Mock) or drops containing spores of WT or ΔBcpnl1 mutant strains of B. cinerea. (A) One day after inoculation, treated leaves were cut in half and each half was immediately colored with Trypan Bleu (TB) for cell death visualisation (on the left of each image) or with 3,3’ - diaminobenzidine (DAB) for oxidative burst visualisation (on the right in each image). (B,C) Zoom of a WT (B) or mutant (C) infected leave colored with TB. Black arrows show the fungus and the red arrow shows a spot of induced necrosis. Scale bar = 50µm. Images were taken with a ZEISS Axio Zoom V16. Scale bars = 500µm. Images were taken with an ZEISS Axio Zoom V16.

### The impact of host nitrogen nutrition on plant JA-related gene expression is partially altered during infection with the ΔBcpnl1 mutants

When plants were infected with the *ΔBcpnl1* mutants, we observed at 1dpi, both at high and low N, a lower response of most tested plant defense-related genes compared to the WT strain B05.10. Indeed, the JA marker gene *AtPDF1*.2a, the chitinase *AtPR4* and *AtPAD3* involved in camalexin biosynthesis were down-regulated at 1 dpi (Figure 8A). Accordingly, the higher expression level of the glucanase *AtPR2* and the transcription factor (TF) *AtWRKY70* observed with the *ΔBcpnl1* mutants correspond to a reduced response since these genes are repressed upon infection. Moreover, *AtJOX3* and the transcription factors *AtJAZ1,3,4* genes were also down-regulated in plants infected with the mutants. Nevertheless, *AtPR1* was not differently expressed between plants infected by the mutant or the WT strain at low N and slightly reduced in two mutants at high N. This result indicated that infection with the *ΔBcpnl1* mutants can still trigger early SA-mediated defense even if the overall plant response to the mutants is limited compared to the WT.

However, it is worth to note that while the host nitrogen nutrition still impacts the level of expression of *AtPR1, AtPR2* and *AtPAD3,* nitrogen nutrition didn’t affect anymore *AtJAZ3,4* and *AtJOX3* expression upon infection with *ΔBcpnl1* mutants (Figure 8B).

Interestingly, by looking at a later stage of the infection process we observed at 2 dpi a strong up-regulation of *AtPDF1.2* during infection with the *ΔBcpnl1* mutants compared to the wild-type indicating a stronger activation of the JA defense pathway (Figure 8C). Furthermore, this up-regulation was even stronger in high N conditions. These results are consistent with the observed incapacity of the mutants to produce the JA repressing high DP unsaturated OGs.

**Figure 8:**
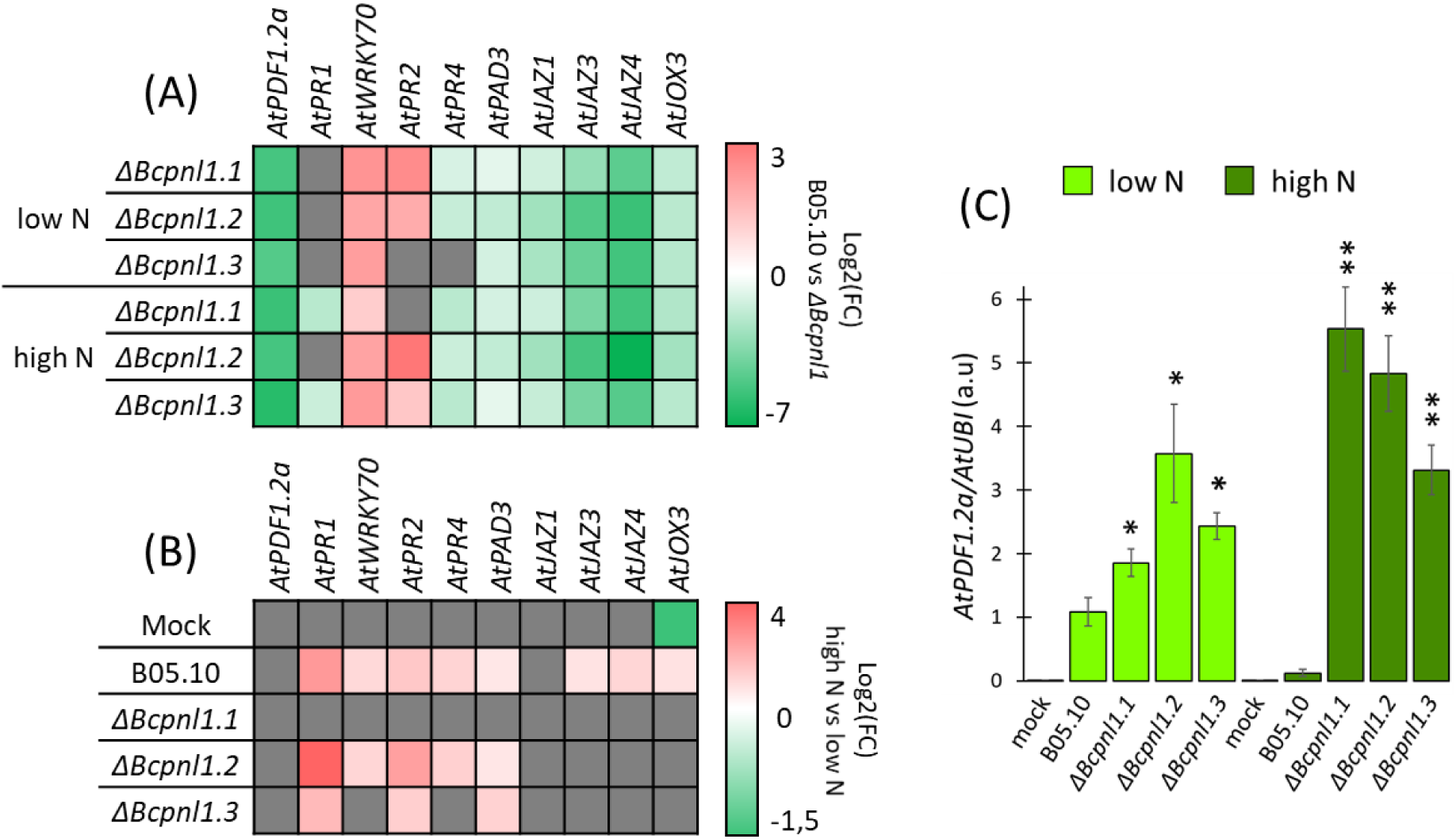
Differences in plant defense gene expression during infection of low and high N A. thaliana plants with the B05.10 reference strain and the ΔBcpnl1 mutant strains. The log2 fold change of transcript accumulation at 1 dpi are represented for defense genes between B05.10 and the mutants (A) and between high N and low N conditions (B). Only significant differences obtained with a two-sample t-test are shown, p<0.05. Unsignificant differences are represented with a grey square. Red: more expressed in B0510, green: more expressed in ΔBcpnl1 mutants. (C) Transcript accumulation of AtPDF1.2a in A. thaliana leaves at 2 dpi with B05.10 and the ΔBcpnl1 mutant strains. In each condition, a two sample t-test was conducted between the B05.10 strain and the mutant strains: *p<0.05,**p<0.01. Plant gene targets were normalized with AtUBI4. Similar results were obtained using AtAPT1 for normalization. Three independent experiments were conducted with similar results.

## Discussion

*Botrytis cinerea*, like all pectin-degrading fungal pathogens, relies on its pectinases to break the cell wall barrier and release nutrients that support fungal growth and development. However, the impact of abiotic factors such as nitrogen nutrition on this cell wall degradation process, as well as the role of PNLs as pathogenicity factors in *B. cinerea* pathogenesis, had never been reported. In this study, we were able to point towards the involvement of BcPNL1 in orchestrating part of the interaction between host nitrate nutrition and disease severity.

We first showed that more PNL products were produced in high N plants infected with the B05.10 wild type strain of *B. cinerea*, and this higher production correlates with a higher upregulation of *AtJOX3* and *AtJAZs.* These genes are involved in repression of the JA pathway which is strongly implicated in efficient defence strategy against *B. cinerea* (AbuQamar et al., 2017; Thomma et al., 1998). This pathway is induced by long methyl esterified PNL products, accumulated upon plant infection with the *ΔBcpme1ΔBcpme2* mutant, hypervirulent on *A. thaliana.* With the *ΔBcpme1ΔBcpme2* mutant, these high DP (>5) PNL products, that cannot be further degraded by PGs induce an up-regulation of genes involved in JA pathway repression such as *AtJOX3* and *AtJAZs* (Voxeur et al., 2019). Therefore, to explain why high DP PNL products were accumulated in high N infected plants, we assessed if the host nitrogen nutrition could impact the transcriptional regulation of *BcPNLs* and/or *BcPMEs.* However, we did not observe any downregulation of these genes in high N condition that would have explained the accumulation of high DP PNLs OGs. Thereby, potential differences in the composition or structure of the plant cell-wall affecting pectin accessibility to theses enzymes might explained the observed differences. To investigate further on the implication of pectin lyase activity in the relationship between nitrogen nutrition and disease susceptibility, we characterized the role of the pectin lyase BcPNL1, encoded by the most expressed *PNL* genes in the early phase of infection (Figure 1B) and from which the *in vitro* PNL activity was previously confirmed (Voxeur et al., 2019).

Thanks to the recently described CRISPR-Cas9-mediated gene replacement in *B. cinerea* (Leisen et al., 2020), we have obtained and then characterized *ΔBcpnl1* mutants (Figure 2). *In vitro*, the mutants presented normal growth with glucose or highly methylesterified pectin as the sole carbon source (Figure S2). Similarly, *Δpnl1* mutants of *Penicillium digitatum* displayed no growth defects *in vitro* (López-Pérez et al., 2015). However, we observed a delayed spore germination of *ΔBcpnl1* mutants on onion epidermis (Figure S2). In the transcriptome analysis of germinating spores on a plant mimicking surface, Leroch et al. (2013) showed that *BcPNL1* expression is strongly induced after only one hour of incubation. These observations suggest that *BcPNL1* might be implicated in early germination events upon perception of the plant surface. Consistently, *BcPNL1* mutation was accompanied with a strong downregulation of the expression of other fungal pectinases during infection (Figure 5).

In other phytopathogens such as *Pectobacterium carotovorum*, *Penicillium digitatum* and *Verticillium dahlia,* PNL activities represent important determinants of pathogenicity(Chen et al., 2016; López-Pérez et al., 2015; Pirhonen et al., 1991). Concerning *B. cinerea*, to our knowledge no study had clearly investigated the role of *BcPNL* genes. *In planta,* we observed a reduced pathogenicity of *ΔBcpnl1* mutants restored in the complemented strains (Figure 4, 5) demonstrating that PNL activity is an important determinant of *B. cinerea* pathogenicity too. Furthermore, the reduced pathogenicity of the *ΔBcpnl1* mutants on different host plants also suggests that this pathogenicity factor is not linked to host specificity. Supporting this hypothesis, Cotoras & Silva (2005) observed in WT *B. cinerea* strains isolated from tomato and grape a similar level of secretion of PNLs. In other fungal pathogens such as *V. dahlia* and *P. digitatum*, mutants of PNLs were also hypovirulent on their respective hosts (Chen et al., 2016; López-Pérez et al., 2015). Authors not only mention their role in promoting penetration through cell wall degradation but also their necrotizing activity. In our study, according to trypan blue colorations, initial necrosis of the plant cells seemed to be reduced in presence of the *ΔBcpnl1* mutants (Figure 5). Movahedi & Heale, (1990) showed that a PNL purified from *B. cinerea* germinating spores had a cell death activity on carrot cell culture and was able to produce elicitors from cell wall components. Since *BcPNL1* was the only *PNL* gene that was strongly expressed in the early interaction, it might be the same enzyme that the authors purified and characterized as a necrosis-inducing protein. Finally, we observed that the increased susceptibility of high N plants to *B. cinerea* was still observed with *ΔBcpnl1* mutants suggesting that BcPNL1 pectinolytic activity is not the only factor determining the impact of host nitrogen nutrition on disease severity.

Consistently, in the *ΔBcpnl1* mutants, pectin degradation capacities were strongly affected both at high and low N. Far less PNL-derived cell wall products were detected upon infection with the *ΔBcpnl1* mutants (Figure 5) which corroborates the absence of functional compensation observed with other pectinases at the transcriptional level. It is known that pectinases are regulated at the transcriptional level by pectin and pectic components (Yadav et al., 2009), thus, specific BcPNL1 products released upon degradation might induce genes coding for enzymes capable to achieve the next degradation step. This could explain the absence of compensation by *BcPMEs* and *BcPGs* of the *BcPNL1* deletion. Overall, deletion of *BcPNL1* seemed to modify deeply the kinetics of pectin degradation and OG turnover during infection which can explain the delayed pathogenicity of the mutants at high and low N.

During host-pathogen interactions, OGs produced by pectinases are important DAMPs which modulate immunity in plant (Davidsson et al., 2017; Pontiggia et al., 2020; Voxeur et al., 2019). In our study, we found a reduced pathogenicity in the presence of the *B. cinerea ΔBcpnl1* mutants on *A. thaliana* leaves at high and low N. At the transcriptional level, we observed at 1 dpi, a reduced expression or repression of all defense-related genes tested (Figure 8) compared to the WT at high and low N. We attributed this result to the delayed germination of *ΔBcpnl1* mutants and to the down regulation of the expression of all pectinase genes. When we compared the expression level of defense-related genes between high and low N plants infected by *ΔBcpnl1* mutants, we observed that, whereas the impact of nitrogen nutrition on defense gene expression was conserved, we did not observe it anymore on JA-supressor genes. Furthermore, at 2 dpi when symptoms are observed both in plants infected with WT and *ΔBcpnl1* mutant strains, a stronger induction of *AtPDF1.2a*, a marker of the JA defense pathway was observed with the *ΔBcpnl1* mutants at both nitrate conditions but with a higher fold change in high N (Figure 8). These data underline on one hand that high DP unsaturated OG produced by BcPNL1 are capable of suppressing plant JA-mediated defense activation and on the other hand that these OGs have a significant role in the nitrogen-dependent down-regulation of plant defense.

To conclude, we found that host nitrogen nutrition impacts the plant pectin sensitivity to the fungal pectin lyase BcPNL1 that results in the production of more high DP PNL products at high N which partially hijack the DAMP-mediated immunity.

## Materials and Methods

### Fungal and plant material

*B. cinerea* strain B05.10 collected from *Vitis* in Germany (Quidde et al., 1998) was used throughout this work as the WT reference strain. Strains of *B. cinerea* were routinely cultured on Potato dextrose agar (PDA, Difco) or Malt Agar (1% malt extract, 1.2% agar) at 23°C under constant light. For fungal growth rate measurements, we used Czapeck minimal media containing 2.55 g/L NaNO_3_, 0.5 g/L KCl, 0.5 g/L MgSO_4_.7H_2_O, 10 mg/L FeSO_4_.7H_2_O, 1 g/L K_2_HPO_4_, 10 mg/L Na_2_MoO_4_.2H_2_O and either 30 g/L glucose or 10 g/L of highly methylesterified citrus pectin (sigma) as the sole carbon source. *A. thaliana* ecotypes Col-0 and Ws were obtained from the INRAE-Versailles collection. For all experiments, plants were cultured in growth chambers (65% relative humidity, 8 hr photoperiod, 21°C) for 6 weeks on nonsterile sand either supplied with a nutrient solution containing 0.5 mM NO_3_^-^ (Low N) or 10 mM (High N). Tomato plants (Monalbo) were cultured in the same conditions for 6 weeks but were only supplied with 10 mM NO_3_^-^ nutritive solution.

### Construction of the ΔBcpnl1 mutants and complemented strains

*ΔBcpnl1* mutants and complemented strains were obtained by using CRISPR-Cas9 mediated homologous recombination (HR) (Figure 2, Figure S3) as described in Leisen et al. (2020). Briefly, fresh spores of the B05.10 strain (1.10^8^ conidia) were cultured overnight in 100 mL of HA liquid medium (1% malt extract, 0.4% glucose, 0.4% yeast extract, pH 5.5). The obtained fungal biomass was washed twice in KCl/NaPi buffer (0.6 M KCl, 100 mM sodium phosphate pH 5.8) and then incubated in KCl/NaPi buffer with 1.35 % (w/v) protoplasting enzymes (Glucanex) 1h at 28° on a rotary shaker at 60 rpm. Protoplast were then washed twice with TMS buffer (1 M sorbitol, 10 mM MOPS, pH 6.3) and concentrated in TMSC buffer (TMS + 50 mM CaCl_2_) at the desired concentration (2.10^7^.100 µL^-1^). PEG-mediated transformation was achieved by addition of PEG solution (0.6 g ml^-1^ PEG 3350, 1 M sorbitol, 10 mM MOPS, pH 6.3) to the protoplasts previously incubated for 10 min on ice with the ribonucloprotein (RNP) complex and the DNA repair template (10 µg). The RNP complex was obtained by incubating for 30 min at 37°C in 15 µL of EngesCas9 specific buffer (NEB), 6 µg of the commercial EngenCas9 (NEB) with 2 µg of the freshly synthetized SgRNA1 obtained with the HiScribe™ T7 High Yield RNA Synthesis Kit (NEB). DNA repair templates containing the fenhexamid resistance cassette surrounded by 60 bp of the target gene for HR were obtained by PCR. For the *ΔBcpnl1* mutants primer X1 and X2 were used on the pTEL-fenh plasmid kindly provided by Pr. Matthias Hahn (Kaiserslautern, Germany). For the complemented strain c1, primers X3 and X4 were used to amplify the coding sequence of *PNL1* together with the native promoter (1,5 kb) and the terminator (0,5 kb). For the complemented strains c2 the oliC promoter and the gluc terminator were amplified from the pndh-ogg plasmid (Schumacher, 2012), kindly provided by Muriel Viaud (INRAE, Saclay), using the primer pairs X5-X6 and X7-X8 respectively whereas the *PNL1* coding sequence was amplify with X9 and X10. To amplify the hygr cassette, the X11 and X12 primers were used on the pndh-ogg. Finally, the two or four fragments respectively were then assembled by overlapping tail PCR using X13 and X14 to form the final fragment carrying the 60 bp for HR in an intergenic region of chromosome 3 as described in Jeblick et al. (2023). The protoplasts were then plated on SH agar medium (0.6 M sucrose, 5 mM Tris-HCl pH 6.5, 1 mM (NH_4_)H_2_PO_4_, 9 g.L^-1^ bacto agar) supplemented with 1 µg.mL^-1^ fenhexamid or 40 µg.mL^-1^ hygromycin B for regeneration. Resistant transformants were then subcultured with the selection pressure on HA agar (HA, 1.5 % agar) and once again on PDA and MA for stock solution of spores and DNA extraction respectively. Finally, correct homokaryotic integration was verified by PCR on genomic DNA extracted as described in Feng et al. (2010). Unique integration of the cassette in the mutants was then controlled by Southern blot using DIG-High Prime DNA Labelling and Detection Starter Kit I (Sigma). Primers used for these experiments are listed in Table S1. Selection of the SgRNA protospacer sequences was done with the online CRISPOR tool (Concordet & Haeussler, 2018).

### Infection experiments

For all infection experiments, spores of each strain were harvested from 2 weeks-old PDA cultures. For pathogenicity assays, RT-qPCR experiments and colorations, detached leaves were inoculated on their upper surface with 20 µL drops containing 5.10^5^ conidia.mL^-1^ in ½ Potato dextrose broth (PDB, Difco). Leaves were then kept in a Petri dish under high humidity conditions and daylight at 23°C. Fungal DNA quantification in infected leaves was performed according to Gachon & Saindrenan (2004). For OG production experiments, triplicates of 10 detached leaves were directly immersed in 35 mL of a *B. cinerea* spores suspension containing 3.10^5^ conidia.mL^-1^ in Gamborg’s B5 basal medium, 2% fructose and 10 mM phosphate buffer pH 6.4. Samples were incubated at 23°C on a rotary shaker at 80 rpm for 15 h or 18 h as described in Voxeur et al. (2019).

### DAB and BT coloration experiments

For coloration assays, leaves were infected as described above and cut in half 24 h after inoculation prior to coloration. Detection of H_2_O_2_ production was achieved by 3,3’-diaminobenzidine (DAB) staining. Briefly, samples were immerged in an aqueous solution containing 1 mg.mL^-1^ DAB (pH 3.6, Sigma) and incubated overnight at RT on a rotary shaker. To discolour the leaves, chlorophyll was extracted by incubation in 75% ethanol. To detect cell death, Lactophenol Trypan Blue (TB, Sigma) was performed as described in Chassot et al. (2008). Pictures were taken on a ZEISS Axio Zoom V16.

### Gene expression analysis

For RT-qPCR, in each condition total RNA was extracted from a pool of 8 leaves and extracted using TRIzol reagent (Invitrogen). Three independent biological replicates were analysed. Reverse transcription was performed using an oligo-dT20 for a primer and Superscript II RNaseH-reverse transcriptase (Invitrogen). PCR was performed on a Applied Biosystems™ QuantStudio™ 5 Real-Time PCR System with SYBR Green PCR MasterMix (Eurogentec). Each reaction was performed on a 1:20 dilution of the cDNA, synthesized as described above, in a total reaction of 5 μL. Gene expression values were normalized to expression of the *A. thaliana AtAPT1* and *AtUBI4* genes or *B. cinerea BcACTA* and *BcUBI* (Ren et al., 2017). We obtained similar results with both genes, therefore results with only one reference gene are shown. Specific primer sets are given in Table S2.

### Analysis of OG production

After 15 h or 18 h of incubation of the leaves in a suspension of spores an equal volume of ethanol 96% was added in order to precipitate the largest molecules. The methodology outlined previously (Voxeur et al., 2019) was followed for the analysis of OG production. In summary, samples were diluted to 1 mg/ml in 50 mM ammonium formate containing 0.1% formic acid. Chromatographic separation utilized the ACQUITY UPLC Protein BEH SEC Column (125 Å, 1.7 μm, 4.6 mm × 300 mm; Waters Corporation, Milford, MA, USA) with elution in 50 mM ammonium formate and 0.1% formic acid at a flow rate of 400 liters/min and a column oven temperature of 40°C. The injection volume was 10 μl. Mass spectrometry detection was conducted in negative mode with the following settings: end plate offset set voltage at 500 V, capillary voltage at 4000 V, nebulizer pressure at 40 psi, dry gas flow at 8 liters/min, and dry temperature at 180°C. Peak annotation was performed based on accurate mass annotation, isotopic pattern, and MS/MS analysis, as described previously (Voxeur et al., 2019).

## Acknowledgments

This work has benefited from the support of IJPB’s Plant Observatory technological platforms.

## Author contributions

Conceptualization: AD, AV, SV, MCS, MF

Methodology: AD, AV, MCS, MF

Investigation: AD, AV

Visualization: AD, AV

Supervision: MF, MCS, AV

Writing—original draft: AD, AV

Writing—review& editing: AD, AV, SV, MCS, MF

