## supplemental material for "Unravelling the Interplay of Nitrogen Nutrition and the *Botrytis cinerea* pectin lyase BcPNL1 in Modulating *Arabidopsis thaliana* Susceptibility"

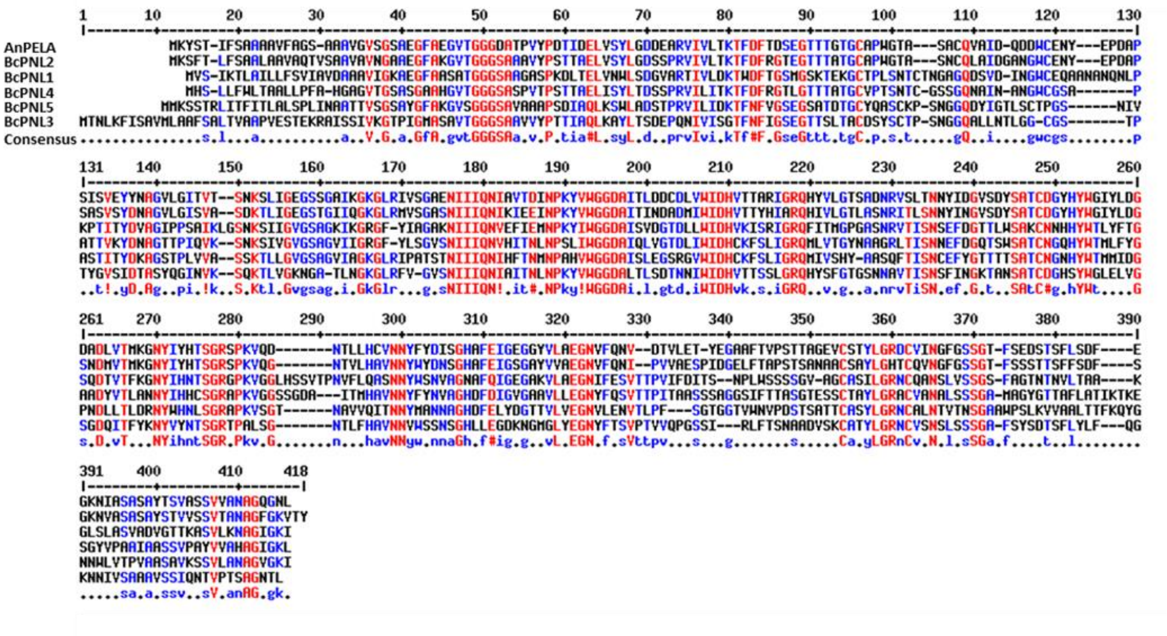


***Figure S1: The pectin lyase protein family in Botrytis cinerea***. *Multiple Sequence Alignment of the Pectin Lyases protein family in B. cinerea and the pectin lyase AnPELA from Aspergillus niger. Red and blue letters indicate conserved amino acids between all or several sequences respectively. Alignment was performed with the MultAlign online software.*

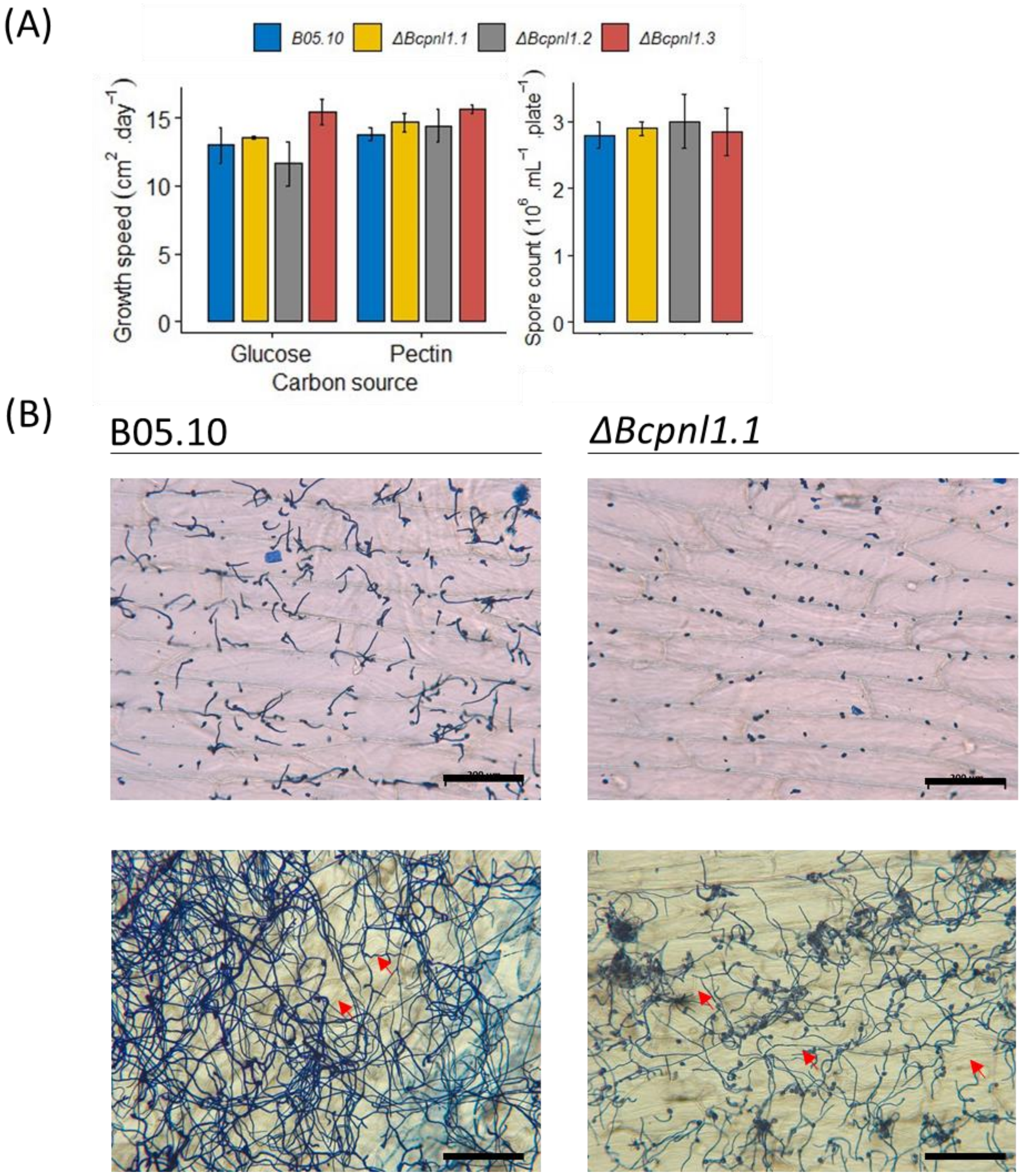


***Figure S2*: *Phenotypic characterization of the ΔBcpnl1 mutants*.** *(A) In vitro linear growth speed with glucose or citrus pectin as carbon source and spore production on PDA medium are compared between B05.10 and the ΔBcpnl1 mutant strains in the left and right charts respectively. Data are the means of triplicates (±SE). In each condition, two sample t-test was conducted between the B05.10 strain and the mutants with no significant differences. (B) Coton blue coloration of the fungus on onion epidermis. Onion epidermis were inoculated with spore droplets of either B05.10 or the ΔBcpnl1 mutant strains in ½ PDB and incubated for 8h (top panel) or 24h (lower panel) before coloration and observation under the microscope. Red arrows show penetrating hyphae. Since results were the same with ΔBcpnl1.1/2/3, only the observations for ΔBcpnl1.1 are presented. Scale bar = 200 µm.*

| | | |
|---|---|---|
| BcPNL2 fwd | 5’-TCGTGTCATCGTTCTCACCA-3’ | RT-qPCR |
| BcPNL2 rev | 5’-TGGAACCCTCACCGATCAAA-3’ | RT-qPCR |
| BcPNL3 fwd | 5’-CAACCAGGCTCTTCGATTCG-3’ | RT-qPCR |
| BcPNL3 rev | 5’-CACAATGTGTTTCCTGCCGA-3’ | RT-qPCR |
| BcPNL4 fwd | 5’-AGCTGGCAGACTCACAATCT-3’ | RT-qPCR |
| BcPNL4 rev | 5’-GGCCTGCGACATTGTAGAAG-3’ | RT-qPCR |
| BcPNL5 fwd | 5’-AGCCGTCTAATGGTGGTCAA-3’ | RT-qPCR |
| BcPNL5 rev | 5’-CCAAACATGTGCAGGGTTCA-3’ | RT-qPCR |
| AtAPT1 fwd | 5’-CCTATTGCGTTGGCTATTG-3’ | RT-qPCR |
| AtAPT1 rev | 5’-TCTTCACTCCTACTCGTTC-3’ | RT-qPCR |
| AtUBI4 fwd | 5’-TGACACCATCGACAACGTGA-3’ | RT-qPCR |
| AtUBI4 rev | 5’-GAGGGTGGACTCCTTCTGGA-3’ | RT-qPCR |
| AtPDF1.2 fwd | 5’-TTTGCTTCCATCATCACCCTTATCTT-3’ | RT-qPCR |
| AtPDF1.2 rev | 5’-ACACTTGTGTGCTGGGAAGA-3’ | RT-qPCR |
| AtPR1 fwd | 5’-CTGGCTATTCTCGATTTTTAATCG-3’ | RT-qPCR |
| AtPR1 rev | 5’-TCCTGCATATGATGCTCCTTATTG-3’ | RT-qPCR |
| AtPAD3 fwd | 5’-CCTCGTCCTCTTACCCCTGA-3’ | RT-qPCR |
| AtPAD3 rev | 5’-GAAGCTTCTTTGGACCCGGA-3’ | RT-qPCR |
| AtWRKY70 fwd | 5’-CCCAAGAAGTTACTTTAGATGCAC -3’ | RT-qPCR |
| AtWRKY70 rev | 5’- TTGCTCTTGGGAGTTTCTGC-3’ | RT-qPCR |
| AtJOX3 fwd | 5’- GAACCAGCTCCTCATGCTTT-3’ | RT-qPCR |
| AtJOX3 rev | 5’- GGGTTCACATCACTCTGTG-3’ | RT-qPCR |
| AtJAZ1 fwd | 5’- GGAAAATAAGGCATGGCTTGG-3’ | RT-qPCR |
| AtJAZ1 rev | 5’- CGTAACCTCTCGATCACATAGC-3’ | RT-qPCR |
| AtJAZ3 fwd | 5’-TTCAGATTTGGTACCTCAGCC -3’ | RT-qPCR |
| AtJAZ3 rev | 5’-GTTGCTTAATTGGTACGGTGC -3’ | RT-qPCR |
| AtJAZ4 fwd | 5’- ACAGTCTCCATGTCCGAACC-3’ | RT-qPCR |
| AtJAZ4 rev | 5’-TTGCTTTTACAAGCCCAGTG -3’ | RT-qPCR |

***Table S1. List of primers used to produce the strains studied in this work.***

| | | |
|---|---|---|
| BcPNL2 fwd | 5’-TCGTGTCATCGTTCTCACCA-3’ | RT-qPCR |
| BcPNL2 rev | 5’-TGGAACCCTCACCGATCAAA-3’ | RT-qPCR |
| BcPNL3 fwd | 5’-CAACCAGGCTCTTCGATTCG-3’ | RT-qPCR |
| BcPNL3 rev | 5’-CACAATGTGTTTCCTGCCGA-3’ | RT-qPCR |
| BcPNL4 fwd | 5’-AGCTGGCAGACTCACAATCT-3’ | RT-qPCR |
| BcPNL4 rev | 5’-GGCCTGCGACATTGTAGAAG-3’ | RT-qPCR |
| BcPNL5 fwd | 5’-AGCCGTCTAATGGTGGTCAA-3’ | RT-qPCR |
| BcPNL5 rev | 5’-CCAAACATGTGCAGGGTTCA-3’ | RT-qPCR |
| AtAPT1 fwd | 5’-CCTATTGCGTTGGCTATTG-3’ | RT-qPCR |
| AtAPT1 rev | 5’-TCTTCACTCCTACTCGTTC-3’ | RT-qPCR |
| AtUBI4 fwd | 5’-TGACACCATCGACAACGTGA-3’ | RT-qPCR |
| AtUBI4 rev | 5’-GAGGGTGGACTCCTTCTGGA-3’ | RT-qPCR |
| AtPDF1.2 fwd | 5’-TTTGCTTCCATCATCACCCTTATCTT-3’ | RT-qPCR |
| AtPDF1.2 rev | 5’-ACACTTGTGTGCTGGGAAGA-3’ | RT-qPCR |
| AtPR1 fwd | 5’-CTGGCTATTCTCGATTTTTAATCG-3’ | RT-qPCR |
| AtPR1 rev | 5’-TCCTGCATATGATGCTCCTTATTG-3’ | RT-qPCR |
| AtPAD3 fwd | 5’-CCTCGTCCTCTTACCCCTGA-3’ | RT-qPCR |
| AtPAD3 rev | 5’-GAAGCTTCTTTGGACCCGGA-3’ | RT-qPCR |
| AtWRKY70 fwd | 5’-CCCAAGAAGTTACTTTAGATGCAC -3’ | RT-qPCR |
| AtWRKY70 rev | 5’- TTGCTCTTGGGAGTTTCTGC-3’ | RT-qPCR |
| AtJOX3 fwd | 5’- GAACCAGCTCCTCATGCTTT-3’ | RT-qPCR |
| AtJOX3 rev | 5’- GGGTTCACATCACTCTGTG-3’ | RT-qPCR |
| AtJAZ1 fwd | 5’- GGAAAATAAGGCATGGCTTGG-3’ | RT-qPCR |
| AtJAZ1 rev | 5’- CGTAACCTCTCGATCACATAGC-3’ | RT-qPCR |
| AtJAZ3 fwd | 5’-TTCAGATTTGGTACCTCAGCC -3’ | RT-qPCR |
| AtJAZ3 rev | 5’-GTTGCTTAATTGGTACGGTGC -3’ | RT-qPCR |
| AtJAZ4 fwd | 5’- ACAGTCTCCATGTCCGAACC-3’ | RT-qPCR |
| AtJAZ4 rev | 5’-TTGCTTTTACAAGCCCAGTG -3’ | RT-qPCR |

*Table S2. List of primers used to for RT-qPCR.*
